# Haemosporidian Blood Parasites in nestling birds of prey in Mongolia

**DOI:** 10.1101/2021.06.26.450033

**Authors:** Hamidreza Attaran, Jing Luo, Wang Bo, Reza Nabavi, Hong-xuan He

## Abstract

Haemosporidians are vector-transmitted intracellular parasites that happen in numerous bird species worldwide and may possibly have important effects for wild bird populations. Studies of haemosporidians most dedicated on Europe and North America, and only newly some study in the Neotropics has been done, where the occurrence and influences of the disease have been less considered and are not understood well. In this study we designed a study in the nestling birds of prey in Mongolia. We sampled blood from 72 raptors at 2 different species and evaluated avian haemosporidian infection by two nested PCR protocol and one Real time PCR protocol. Sequencing a portion of the cytochrome b (cyt b) gene of the parasite. From the sampled birds, 10 % were infected by Plasmodium. Inclusive, our findings advocate a high haemosporidian species richness in the bird community of Mongolia. In view of the frequency of local habitat loss that in this area is living, recognize how avian haemosporidians affect bird populations it is very important; in addition, more exhaustive sampling is required to fully understand the range of avian haemosporidian infection in this area.

## Introduction

Understanding the threats of wild animal infectious disease for wildlife conservation is a principal issue [1, 2].This issue is becoming more important as climate change-induced territory changes are rapidly affecting the spreading of parasites and hosts [3]. In the last few decades we had observed an increase in the incidence of some infectious diseases in wild populations [4–6] and these be able to have a significant impression on wildlife populations[1].

Avian haemosporidian parasites (phylum Apicomplexa, order Heamosporida) are belonging to families Heamoproteidae, Plasmodiidae and Leucocytozidae. They are vector-borne parasites and infect most bird families [7]. Heamosporidian parasites are transmited by 17 genera of blood-sucking insects like, biting midges, louse flies, black flies and mosquitoes [7, 8]. The study of these parasites helps us as model of the host-parasite interactions in ecology, evolution and conservation biology. Avian haemosporidian parasites are cosmopolitan in distribution and indirectly reduce host fitness by decreasing their reproductive success and increasing breeding energy cost [9].Therefore, haemosporidian parasites constitutes a selective force on bird population. Accordingly, interactions of haemosporidian parasites and birds have become a model of host-parasite relationship in ecology, conservation and vertebrate management [8, 10]. Among haemosporidian parasites, *Plasmodium* has been linked to the mortality and population-level declines in native birds in some region [11]. Despite of their broad host and geographic distribution, information on haemosporidian pathogenicity is almost based on laboratory experiments with domestic birds [7].

Raptor birds (Accipitriformes and Strigiformes) are located at the top of the food chain. Despite they play important roles in the ecosystem but most studies of avian haemosporidian have been conducted on passerines (Passeriforms), with very little focus on raptor hosts [12, 13].More studies are critical to progress our knowledge of the genetic variability, true diversity and host specificity of blood parasites in raptor birds [14, 15].

Studies on raptors in comparison with passerines are infrequent and have been mostly performed on captive, injured or migratory birds [13, 16–19]. However has been unpaid to the breeding populations of sedentary raptor species, although studies on nestling have several advantages: they are immunologically naive and highly susceptible to infection, detected parasites supposedly originates from the study site, and the acute phase of infection has no effect on sampling since they are immobile [20–25].

Furthermore, while the blood parasites are generally thought to be harmless, we have evidence that infection can be harmful [26]. Studies have suggested that avian haemosporidian infection can cause reduced strength in flight; reduce speed, poor appetite, anemia, weight loss, airsacculitis, arthritis, lower reproductive success, reduced lifespan and death [6, 26–29].

However the environment modulates infection risk. Avian haemosporidian infection has been linked with environmental variables such as altitude, temperature, and precipitation [30].

On the other hand, studies of haemosporidians most dedicated on Europe and North America, and only newly some studies in the Neotropics has been done [31–34].

Presented haemosporidians in isolated areas may pose a threat to naïve endemic birds that have not evolved ways to counteract the infection. For instance, haemosporidian vectors were accidentally introduced to the islands after some Hawaiian birds that suffered large population declines [35, 36]

Mongolia is located in north of China and we don’t have any information about rate of haemosporidian parasites in birds and specially birds of prey there. Because of low human population and huge grass land any information from diseases will be very important specially for china because they are close and any infection can easily come to china from Mongolia.

In the last two decades, molecular biological tools have been obtained to study haemosporidian parasites of bird [37–39] DNA-based techniques make detection of haemosporidians easier, particularly during chronic infections and at early stages of infections, when parasitaemia is low and parasites can be over- looked in blood smears [40]. The progress in this field is directly joined to the development of a standard nested PCR protocol for amplifying a segment of the haemosporidian cytochrome b gene [38, 41, 42]

The goal of this survey was to conduct an initial study of the incidence of avian haemosporidian parasites in the endemic bird area of the Mongolia, home to a variety of biomes and one of the world’s important avian centers of endemism. In addition to habitat-related threats, the local avifauna could also be exposed to infectious diseases, which could exacerbate the effects of habitat reduction on the local populations.

## Material and Methods

### Study area and Sampling

Seventy-two nestling of raptors (70 Saker falcons, and 2 common buzzards) between two till five weeks old were sampled in Mongolia (Bayan district near Ulaan Baatar) during June 2016.

Blood samples after separating serum stored at −80

DNA extraction, PCR amplification, sequencing and parasite detection

Total DNA was extracted from RBC cells using DNA extraction tissue Kit. The existence and quality of DNA was inspected by the spectrophotometer NanoDrop ND-1000.

Samples were screened for blood parasites of the genera Haemoproteus, Plasmodium and Leucocytozoon with two nested PCR protocols and one Real Time PCR Protocol.

### The first nested PCR protocol

For the first way, two modified nested PCR were used to amplify fragments of the cyt b gene (Table 2). The protocol for Haemoproteus/Plasmodium was based on the standard protocol of Waldenström et al. [42]. But with newly designed forward primers, H332F and H350F [43](Fig. 1, Table 2), which match more closely with available GenBank sequences. The protocol produces a 477 bp fragment, which is only one base pair shorter than the fragment produced by the Waldenström et al. protocol [42]. The Leucocytozoon protocol uses the initial primer sets described by Hellgren et al.[41] but with newly designed nested primers (Fig. 1, Table 2). This new protocol produces a 526 bp fragment that encompasses the 478 bp fragment produced by the Hellgren protocol [43].

The second protocol involved a first pre amplification PCR step with the primers Plas1F [44] and HaemNR3 [41], followed by a nested PCR step with the internal primers 3760F [37] and the HaemJR4 [14]. This protocol uses the same PCR conditions as in Waldenström et al. (2004)[42], but it amplifies a 542 bp fragment of the parasite’s cytochrome b that includes the fragment amplified by the primers of Waldenström et al. (2004), and amplifies DNA from all three parasite genera in a single nested PCR.

PCR products were checked in 2% agarose gels stained with GelRed™ (Biotium, USA) under UV light, looking for bands of the appropriate size. A positive control (using DNA template from an infected bird) and eight negative controls (using distilled water instead of template DNA) were included in every 96-well PCR batch to test for reaction performance and to control for PCR contamination (no negative control yielded a positive result).

Also one real time PCR protocol

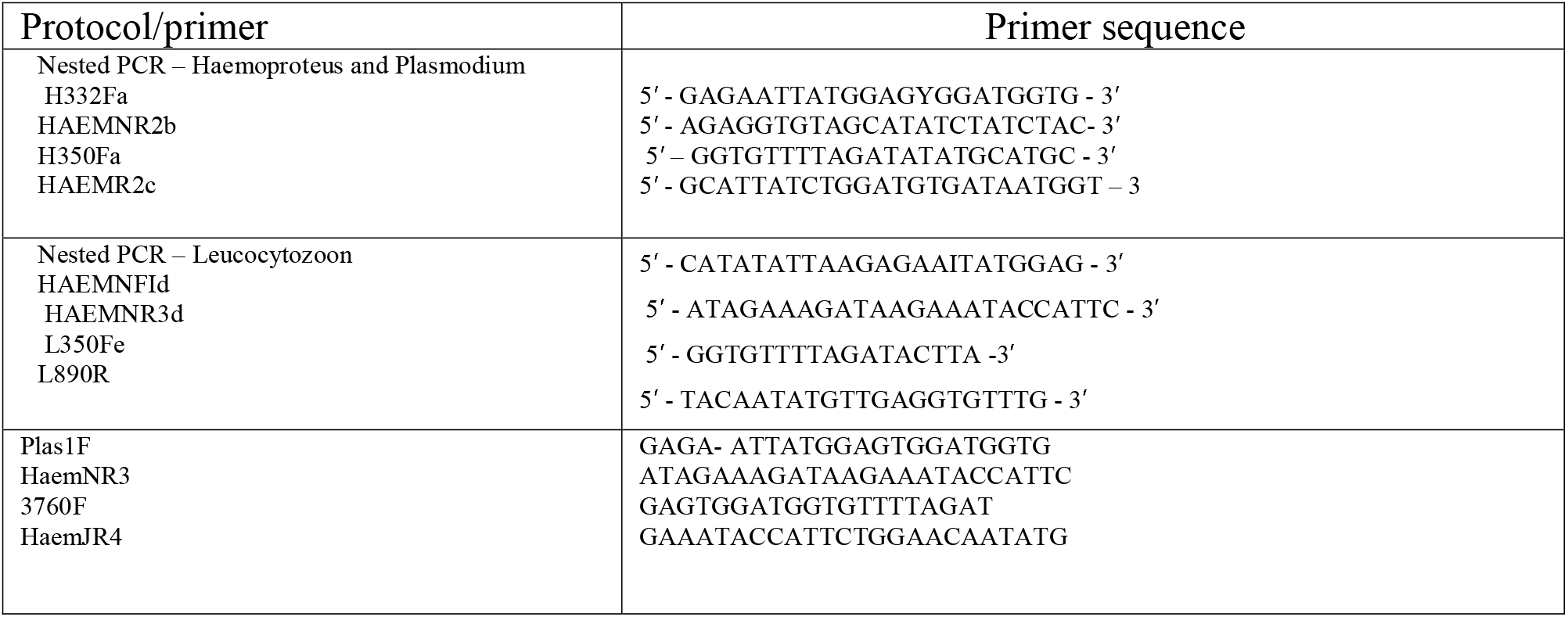
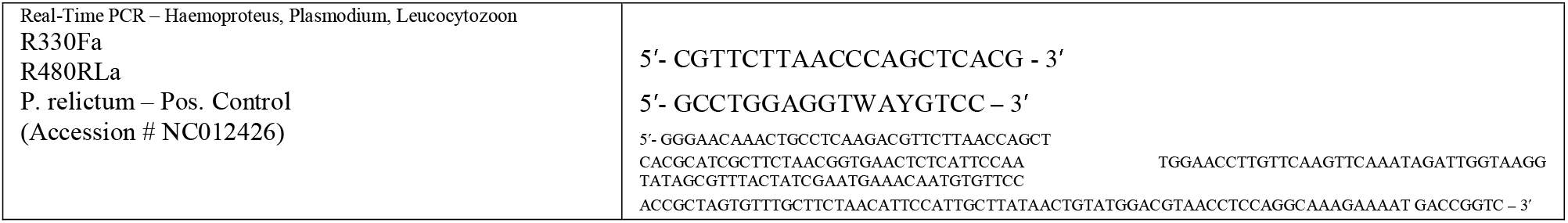

### Phylogenetic analysis

The phylogenetic tree was build using Maximum Likelihood (ML) analysis based on the Kimura 2-parameter model (Kimura 1980) in the program MEGA 7 (Tamura et al. 2013).

## Results

Sequencing of the 7 PCR products that were positive for haemosporidian infection revealed a total of 2 unique sequences that grouped into lineages (Table 2, Figure 2). All sequences identified had a sequence identity of at least 99% to sequences reported in GenBank (Table 2), 2 of these had not been previously reported and have been deposited in GenBank. Four lineages had 100% match to sequences previously reported. One lineage had 100% match to Plasmodium sp., a lineage that has previously been reported in China (Table 2). The phylogenetic analysis of the identified sequences revealed 2 lineages of Plasmodium (Fig. 1). The tree had well-supported nodes, except for the node separating (Fig. 2).

| Sample | Nested PCR (protocol 1) | Nested PCR (protocol 2) | Real-time PCR |
| --- | --- | --- | --- |
| 2 |  | positive |  |
| 6 | positive |  | Positive |
| 18 | positive |  |  |
| 22 |  | positive |  |
| 27 |  | positive |  |
| 34 |  | positive |  |
| 46 |  | positive |  |

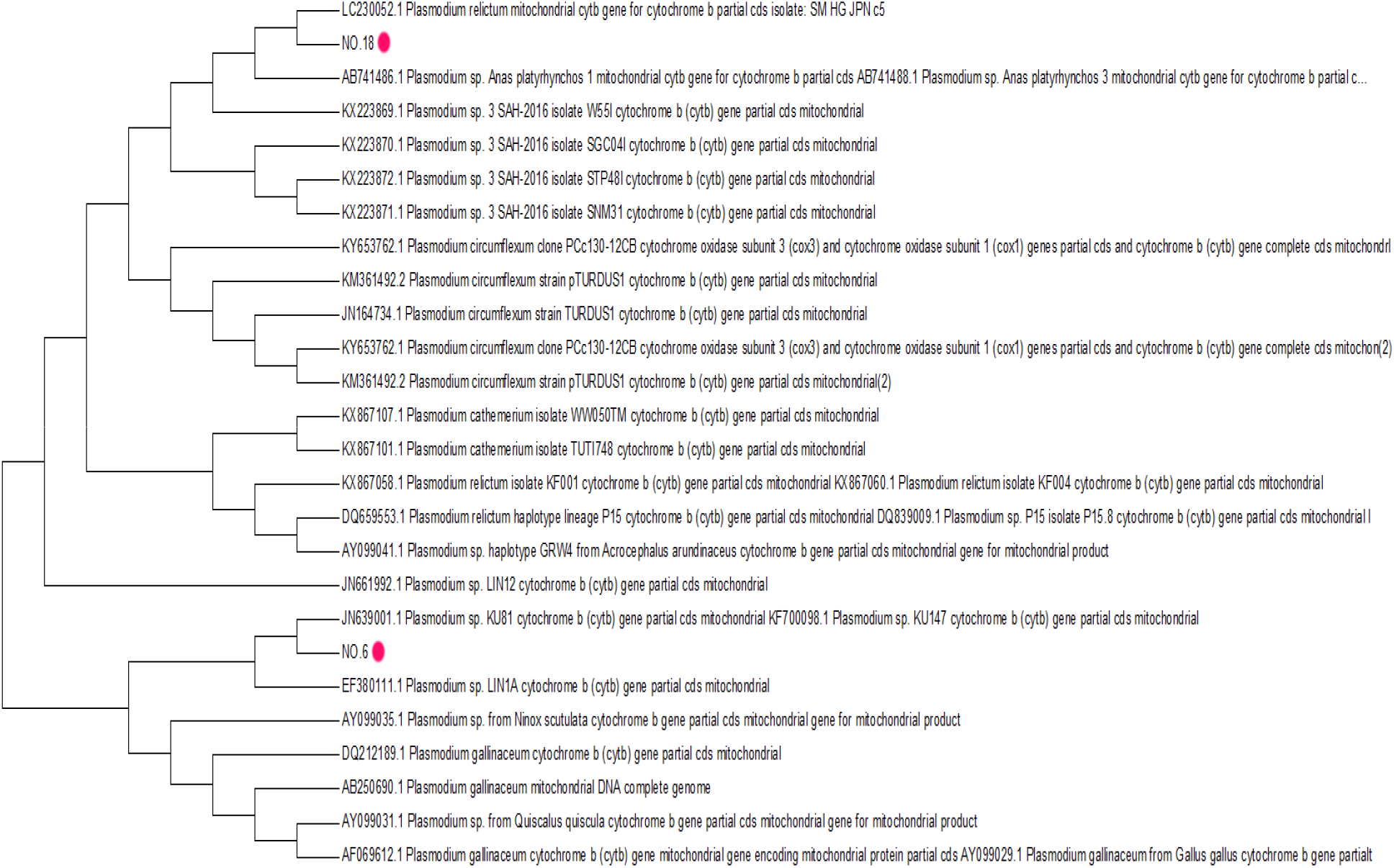

## Discussion

In this study we evaluated infection by haemosporidians in nestling birds of prey from the Mongolia, an important area, and report a possible new blood parasite-host association. Our results obviously showed that the real number of haemosporidian lineages in this area is possible to be much higher than the numbers we found. Considering the protection concerns and its endemic species, the ecology of these parasite - host associations needs to be further discovered.

We found an overall incidence of haemosporidians of 10%. This result concurs with prevalence reported for the González et al. 2015, Krone et al. 2008 and Remple 2004, lierz 2008 [13, 26, 33]. In contrast, a previous large-scale study [22, 45, 46].

In the present study we assessed the molecular identity of avian haemosporidians infecting Raptors, as well as the differences in parasite diversity by three protocols. Although prevalence was generally low.

The purpose of any new screening method is to afford an accurate estimate of parasite prevalence and to offer advantages over already established methods. Although in some previous studies reputed, the real-time PCR protocol as effective as the two most widely used molecular screening methods for haemosporidian parasites in birds [42, 47], in another study reported that all three methods likely leave a small proportion of samples undetected [43]

Recent studies have shown in regions with high avian diversity and in specific host populations [48], the Leucocytozoon diversity would be high[49] accessibility of a screening method that can amplify all three genera can serve up in understanding the accurate diversity and ecology of all three genera of avian haemosporidian parasites. Since now, the only screening process that could distinguish all three genera in a single method were microscopy and the restriction digestion protocol of Beadell & Fleischer [50], but both take drastically more time than the real-time PCR and nested PCR protocols. Although Plasmodium spp. was only found in Mongolia, further studies are needed through a larger elevation gradient to further support our findings. We did not measure other environmental variables that could potentially affect the prevalence of haemosporidians in this area.

Our data revealed incidence of avian Plasmodium in Mongolia from birds of prey. Given Mongolia is considered a center of endemism, high prevalence and diversity of haemosporidians suggests a threat for wildlife.

Once a comprehensive screening of haemosporidians in the wild avifauna is performed, further studies should focus on determining the factors that modulate infection and transmission in the area and assessing the effects of haemosporidian infection on various aspects of host fitness.

---

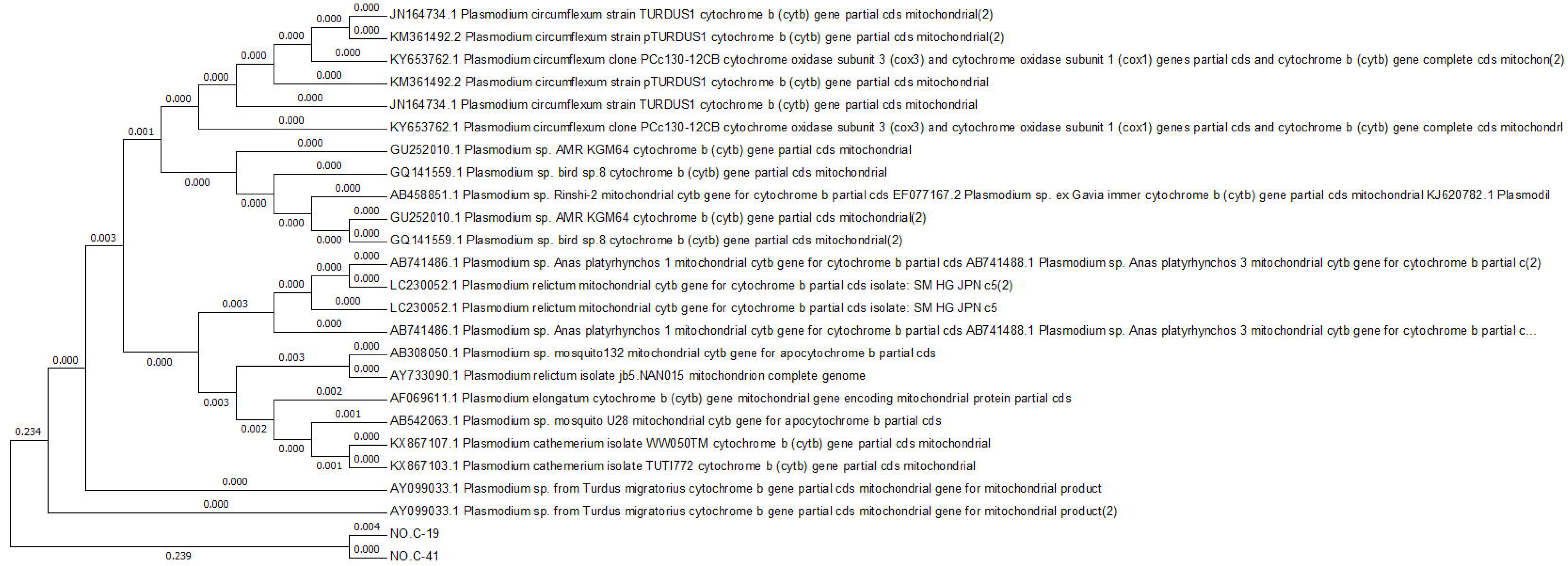

